# TCR germline diversity reveals evidence of natural selection on variable and joining alpha chain genes

**DOI:** 10.1101/2025.08.20.671277

**Authors:** Sreekar Mantena, Ali Akbari, Soumya Raychaudhuri

## Abstract

T cell receptors (TCRs) orchestrate adaptive immunity, yet the complex, repetitive architecture of the TCR loci has impeded systematic characterization of human genetic variation in the genes encoding the TCR. Using public long-read sequencing data from 2,668 donors, we build a near-complete map of common alleles in TCR V, D, and J genes, revealing amino acid variation at almost every position within V genes. We discover pervasive evidence of natural selection on TCR genes, including balancing selection on a TRAJ gene recognizing an immunodominant influenza epitope and positive selection on a TRAV gene. We find TCR allelic polymorphism alters core functional properties of T cells, including thymic fate commitment, phenotypes in diseased tissues, and cell-surface receptor abundance. Collectively, our findings position inherited variation in TCR genes as a key axis of immunological diversity that may shape interindividual differences in immune responses.

## Main Text

Few regions in the human genome have been as profoundly shaped by evolutionary forces as the HLA locus^1,2^. Over millennia, pathogen-driven selective pressures have sculpted extraordinary levels of genetic polymorphism in HLA class I and class II genes. This variation underlies interindividual differences in immune responses, with the HLA locus harboring more GWAS associations than any other genomic region^3,4^.

However, antigen presentation is only one step of the adaptive immune response. The activation of T cells depends on the recognition of antigen–HLA complexes by the T cell receptor (TCR). While the complementarity-determining region (CDR) 3 region of a TCR chain is partially generated through somatic recombination, most of the TCR sequence is encoded by germline V (variable), D (diversity), and J (joining) gene segments. The V gene provides the framework regions that form the structural backbone of the TCR and encodes the CDR1 and CDR2 which contact antigen-HLA complexes, while the D and J genes both contribute to the CDR3 sequence^5^. Many T cell responses are dominated by clonotypes sharing the same germline-encoded V or J gene, with residues in these genes determining key contact residues and binding geometry^5–9^. Thus, germline genetic variation in TCR genes could be an important target of natural selection and may shape interindividual differences in immune responses.

The loci encoding TCR chains, including the TCRα/δ locus (chr14), the TCRβ locus (chr7), and the TCRγ locus (chr7), have a complex genetic architecture marked by extensive gene duplications with high sequence homology among paralogs. For example, the TCRβ locus encodes 67 Vβ genes, several of which are identical or nearly identical in sequence^10^. Short-read technologies are stymied by multi-mapping and are incapable of accurately surveying polymorphism in the TCR loci and other highly repetitive genomic loci^11–15^. Thus, TCR loci remain amongst the least characterized regions of the human genome; the largest studies to date survey fewer than 50 individuals, leaving human genetic diversity in TCR genes poorly understood^16–19^. Recent advancements in long-read sequencing technologies present a unique opportunity to faithfully characterize human genetic polymorphism in these key immune loci^16,19^.

Here, we leverage long-read sequencing data from 2,668 donors to reveal extensive human genetic diversity in TCR V, D, and J genes. We discover this polymorphism has been shaped by natural selection and influences core functional phenotypes of T cells, establishing inherited variation in TCR genes as a meaningful axis of immunological diversity (**Fig. 1a**).

**Figure 1:**
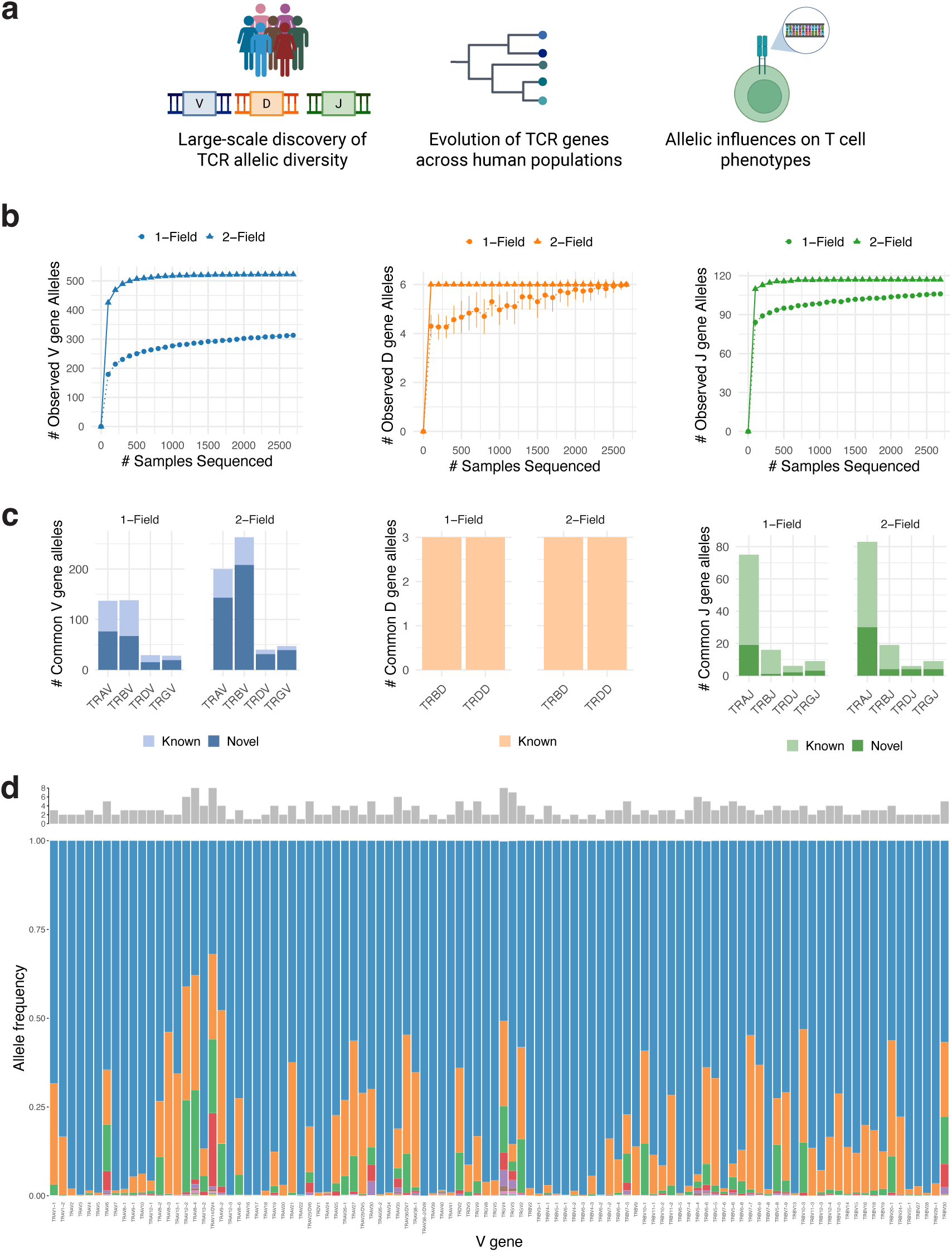
Uncovering the allelic diversity of human TCR genes. **(a)** Schematic of study design. **(b)** Number of common V (left), D (center), and J (right) gene alleles observed as a function of the number of individuals sequenced. **(c)** Total number of known and novel common alleles for TCR V, D, and J gene groups. **(d)** One-field allelic diversity of TCR V genes. Top: Number of common alleles observed per gene. Bottom: Frequency distribution of one-field alleles for each gene; each color denotes a distinct allele.

## Results

### Population-scale discovery of TCR genetic diversity

To discover allelic polymorphisms in TCR loci, we harnessed PacBio HiFi sequencing data from 2,441 individuals from the NIH All of Us CDRv8 release and 227 individuals from the Human PanGenome Reference Consortium release 2^20,21^. We focused on discovering V, D, and J gene alleles, excluding pseudogenes incapable of contributing to functional TCR chains. Following International Immunogenetics Information System^22^ (IMGT) convention, we defined alleles at one-field resolution based on nucleotide changes in the core coding regions (IMGT-REGION). We also defined TCR alleles at two-field resolution based on nucleotide changes in the regulatory regions of TCR genes, as this variation is known to alter TCR gene usage and functionality^23–26^.

At one-field resolution, we observed 664 total V gene alleles, 6 total D gene alleles, and 165 total J gene alleles within our cohort (**Methods; Supplementary Fig. 1a**). Of these, 517 V gene alleles and 84 J gene alleles were novel – representing a 352% increase in the known number of V gene alleles and a 104% increase in the number of J gene alleles (**Supplementary Fig. 1b**). We specifically focused on characterizing common alleles, which we defined as being present at an allele frequency of at least 0.01 in at least one of the five ancestral populations. At one-field resolution, 313 V gene alleles, 6 D gene alleles, and 106 J gene alleles were common (**Fig. 1b-c**). These common V, D, and J gene alleles were directly supported by an average of 9015, 37208, and 33691 reads, respectively (**Supplementary Fig. 1c**). To validate alleles in independent data, we collected publicly-available single-cell TCR sequencing data and found 93% of V and J gene alleles common in donors of European ancestry could be observed in human repertoires (**Supplementary Note 1; Supplementary Fig. 2**). Rarefaction curves indicate our analysis captures over 94% of common TCR V, D, and J gene alleles in each of the five major ancestry groups (**Fig. 1b**; **Supplementary Fig. 3a-c**), enabling us to perform the first population-scale interrogation of the genetic architecture of the human TCR loci.

First, we sought to characterize the broad diversity of TCR gene classes. The 102 TCR V genes had an average of 3.07 common one-field alleles, with some genes harboring up to eight, while the 73 J genes had an average of 1.45 common one-field alleles (**Fig. 1d**). We quantified the variability of each gene across the population with the Shannon entropy of its one-field allele frequencies. TCR V genes were more entropic than J genes, and TRAV genes specifically exhibited greater entropy than TRBV genes (**Supplementary Fig. 3d**). The TRA, TRB, and TRG loci exhibited SNP densities 1.67-, 1.46-, and 2.25-fold higher, respectively, than those of the chromosomes on which they reside, highlighting their unusually elevated polymorphism (**Supplementary Fig. 3e**). Donors of African ancestry exhibited the highest levels of diversity, consistent with previous observations in other genomic regions (**Supplementary Fig. 3**)^27^.

### Presence of loss-of-function alleles

The TCR loci contain dozens of gene segments encoding the same functional unit; this redundancy buffers against the impact of individual loss-of-function (LoF) mutations, allowing nonfunctional alleles to persist at appreciable frequencies. Indeed, some TCR genes are nonfunctional in all humans, while others harbor LoF alleles that vary across individuals. To systematically understand this heterogeneity, we annotated the functionality of TCR alleles, focusing on genes with variable functionality across the population. 20% of TRAV and 22.9% of TRBV genes harbored a common nonfunctional allele, while only 2.0% of TRAJ genes and a 7.7% TRBJ genes did (**Fig. 2a**). Including rare alleles, 57.8% and 54.2% of TRAV and TRBV genes harbored a nonfunctional allele, while 15.7% and 30.8% of TRAJ and TRBJ genes did (**Fig. 2a**).

**Figure 2:**
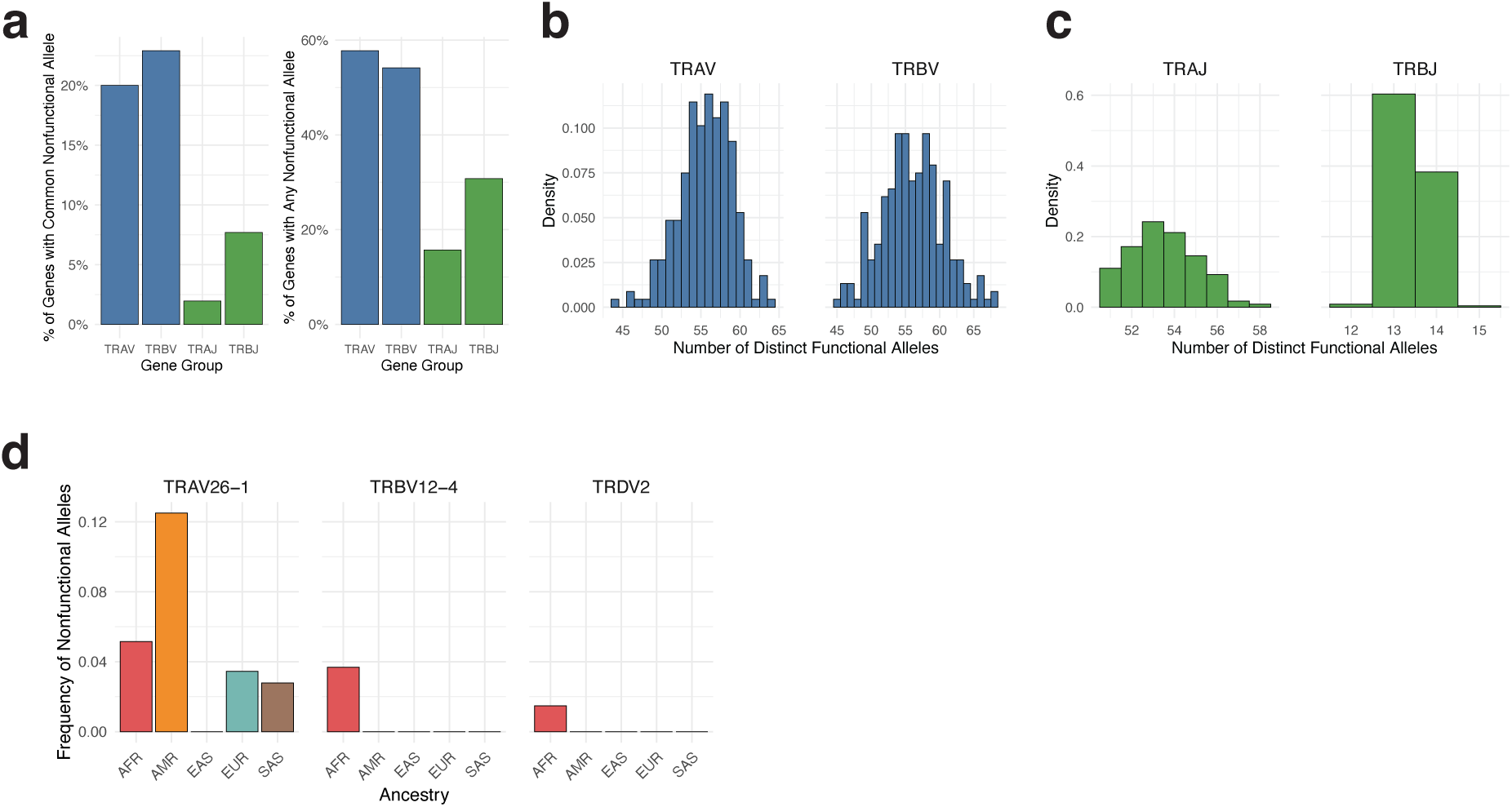
Loss-of-function alleles across TCR V and J genes. (a) Percentage of TCR genes in each gene group that have a common loss-of-function allele (left), and percentage of genes that have a common or rare loss-of-function allele (right). (b) Distribution of the number of distinct functional alleles of TRAV (left) and TRBV (right) genes across donors. (c) Analogous to (b), but for TRAJ and TRBJ genes. (d) Frequency of common nonfunctional alleles in disease-associated TCR genes.

Several V genes, including TRBV10-1, TRAV26-2, and TRBV16, carried a high burden of nonfunctional alleles, with LoF allele frequencies over 0.1. (**Supplementary Fig. 4a**). A subset of genes we discover to have common LoF alleles play key roles in disease contexts (**Fig. 2d**). For example, we identify a LoF allele in TRAV26-1 at a frequency of 0.12 in donors of admixed American ancestry. TRAV26-1 is the dominant Vα gene employed by pathogenic T cells that recognize the HLA-DQ2.5–gliadin-α2 peptide–MHC complex in celiac disease, with germline-encoded TRAV26-1 framework motifs facilitating binding to the gliadin autoantigen^28–30^. In our cohort alone, several donors are homozygous for this LoF allele; they may be protected against developing celiac disease. Furthermore, we find a LoF allele of TRDV2 at 0.02 frequency in donors of African ancestry. TRDV2/TRGV9^+^ cells are the most abundant γδ subset in peripheral blood and play critical roles in anti-tumor and anti-pathogen responses; homozygosity for this LoF allele may impair these responses^31–33^. We also identify a TRBV12-4 LoF allele at 0.04 frequency in donors of African ancestry; the bacterial superantigens SpeC and TSST-1 specifically activate TRBV12-3/12-4^+^ T cells, so carriers of this allele may be partially protected from superantigen-induced cytokine storm^34^.

Surprisingly, we also identified LoF alleles within TCR genes required for the development of unconventional T cell subsets that recognize non-HLA ligands. Specifically, TRAV10, TRBV4-1, and TRAJ33 —required for CD1d-restricted iNKT cells, CD1b/c-restricted T cells, and the predominant population of MR1-restricted MAIT cells, respectively^35–38^ — harbored LoF alleles that were rare in donors of African and admixed American ancestry. The roles of these unconventional subsets remain incompletely understood; studying the phenotypes of individuals genetically unable to generate them offers a natural model for probing their *in vivo* function^39,40^. Furthermore, the presence of these LoF alleles in presumably immunocompetent individuals suggests loss of these T cell subsets may be tolerated.

### Allelic diversity within individuals

Next, we asked if individuals vary in the overall number of distinct functional TCR alleles they carry at one-field resolution. We discovered substantial interindividual variation; donors carried between 44 and 64 (95% empirical CI: 49-61) functional TRAV alleles and between 45 and 68 functional TRBV alleles (95% empirical CI: 47-66). Thus, some individuals have only 66% as many functional TRBV alleles as others (**Fig. 2b-c**). Donors of African ancestry carried, on average, 2.9 more functional TRAV and 5.3 more functional TRBV alleles than donors of European ancestry (**Supplemental Fig. 4b**). To identify the sources of this variability, we used a linear model to partition the variance into three components: LoF allele burden, functional heterozygosity (the number of genes at which individuals are heterozygous for two functional alleles), and genomic deletions. Functional heterozygosity accounted for nearly all of the interindividual variance in functional allele counts (99% for TRAV and 95% for TRBV; **Supplemental Fig. 4c**). To mount an effective response, thymocytes must respond to a vast array of antigens. Heterozygosity at classical HLA genes, which expands the range of peptides presented to T cells, is associated with reduced cancer risk, improved immunotherapy outcomes, and better control of infections like HIV^41–44^. Similar effects may extend to individuals carrying more functional TCR genes, and HLA and TCR heterozygosity may interact to alter immune responses in these disease contexts. Further studies are needed to elucidate these possibilities.

### Natural selection on TCR genes

Having characterized the rich diversity of coding and LoF alleles across human TCR genes, we next asked whether this class of genetic diversity is biologically consequential. Natural selection offers a powerful lens to address this question; evidence of selection on TCR genes would imply that TCR polymorphisms have shaped immune responses to a degree substantial enough to alter organismal fitness. We hypothesized TCR genes may be subject to three distinct selective pressures: (1) Diversifying selection, wherein specific codons across a gene group (e.g., across TRAV genes) evolve under pressure to recognize diverse ligands, (2) Balancing selection, wherein multiple alleles of a specific TCR gene are maintained at intermediate frequency to recognize evolving pathogenic threats, and (3) Positive selection, wherein an particular allele of a specific TCR gene confers a fitness advantage and rises in frequency.

We first tested for diversifying selection; as T cells must recognize broad array of peptide–MHC (pMHC) molecules, selection may have favored the accumulation of amino acid diversity at specific sites within TCR V genes. We constructed dendrograms of common functional TRAV and TRBV genes; as expected, alleles of each gene clustered together (**Fig. 3a-b**). We applied Fast Unconstrained Bayesian Approximation (FUBAR), which uses a Markov chain Monte Carlo framework to estimate per-codon synonymous and nonsynonymous substitution rates to identify sites that have undergone pervasive diversifying selection^45^. Diversifying selection was observed at four codons within the CDRs: IMGT positions 58 and 64 in the CDR2 of TRBV genes, IMGT position 66 in the CDR2/framework region 3 boundary of TRBV genes, and IMGT position 30 in the CDR1 of TRAV genes (**Fig. 3c-d**). Fixed Effects Likelihood (FEL), a complementary maximum likelihood-based approach^46^, confirmed these sites had undergone diversifying selection and also identified IMGT position 38 in the CDR1 of TRBV genes (**Supplementary Fig. 5a-b**). In many crystal structures, the CDR1/2 regions of TCRs contact the helices of HLA molecules^47^. The HLA locus is inherited independently from TCR genes, and CDR1/2-binding regions of HLA helices are not fully conserved^48,49^. Diversifying selection may have acted on germline-encoded CDRs to ensure compatibility with the wide range of HLA alleles an individual can inherit. Additionally, as CDR1/2 often directly contact the peptide^5,8^, antigen-driven pressures may have contributed to diversifying selection at these CDR codons.

**Figure 3:**
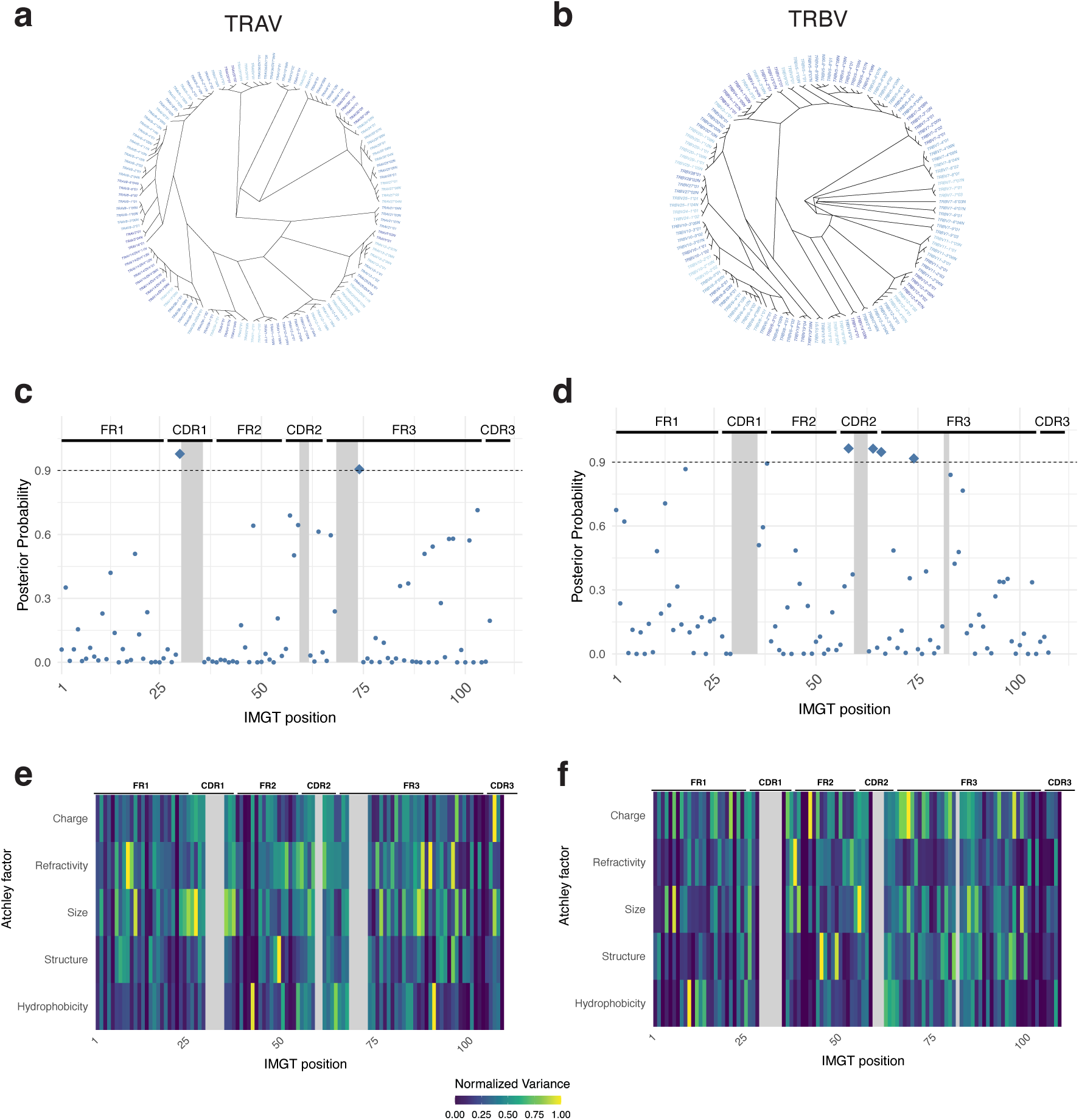
Codon-level diversifying selection on TCR V genes. (a) Dendrogram for common functional TRAV gene alleles. Novel alleles are designated with an ‘N’ suffix. (b) Analogous to (a), but for common functional TRBV gene alleles. (c) Posterior probability of diversifying selection on specific codons in TRAV genes, as determined by FUBAR analysis. Points above the dotted grey line represent significant hits for diversifying selection. Grey bars represent gapped positions. (d) Analogous to (c), but for TRBV genes. (e) Variance in amino acid physicochemical characteristics across common functional TRAV genes. Atchley factors were used to numerically represent physicochemical characteristics, and variance was minimax normalized for each Atchley factor. Columns in grey represent gapped positions. (f) Analogous to (e), but for TRBV genes.

While framework regions are typically considered relatively invariant structural backbones of the TCR, we discovered diversifying selection at IMGT position 74 in the framework 3 region (**Fig. 3c-d**). More broadly, many framework codons in common functional V alleles exhibited high amino acid sequence entropy (**Supplementary Fig. 5c-d**). To determine whether framework diversity reflected substitutions among chemically similar amino acids or functionally distinct ones, we used Atchley factors^50^ to numerically represent physicochemical characteristics. Several framework codons exhibited surprisingly high variance in these physicochemical traits (**Fig. 3e-f**). For example, among all sites, including those in the CDRs, IMGT position 91 in the framework 3 region of TRAV genes exhibited the greatest variance in amino acid hydrophobicity and IMGT position 69 in the framework 3 region of TRBV genes exhibited the greatest variance in amino acid charge. Recent TCR engineering studies have revealed mutations in framework regions modulate CDR loop orientation and flexibility, altering antigen recognition^51–53^.

Moreover, artificial mutations in framework regions influence TCR heterodimer stability and surface expression^54^. Thus, genetic polymorphism in the framework regions likely serves as another source of functional diversity in the TCR repertoire; T cells with identical CDR sequences may differ in antigen sensitivity due to inherited framework variation.

Next, we asked if polymorphisms in TCR genes may be maintained by balancing selection. Classical HLA genes, which present peptides to TCRs, are canonical examples of balancing selection^1,55^. Shifting spatial and temporal pathogen pressures, together with heterozygote advantage, have maintained multiple HLA alleles at intermediate frequencies across human populations^1,56^. To test for balancing selection in TCR genes, we applied BetaScan^57^ across the TCR loci, the HLA locus, and to the TRBorphon locus on chromosome 9, which encodes nonfunctional genes with TRBV-like sequence content and serves as a neutrally evolving control. A hallmark of balancing selection is the build-up of linked polymorphisms whose frequencies mirror that of the selected allele; BetaScan quantifies this clustering of frequency-matched variants to discover balancing selection^57,58^. We found the strongest signal of balancing selection in TRAJ37 (**Fig. 4a**). This signal was robust across all five ancestries, consistent with a pattern of long-term balancing selection (**Supplementary Fig. 6a**). To corroborate this finding, we also computed Tajima’s D, which detects an excess of intermediate-frequency alleles by contrasting mean pairwise diversity with the number of segregating sites^59^. TRAJ37 exhibited markedly elevated Tajima’s D values, exceeding 3.2 in every population (**Fig. 4b**). Consistent with this pattern, the Shannon entropy of TRAJ37 ranked among the top 5% of all TRAJ genes in all ancestries (**Fig. 4c**).

**Figure 4:**
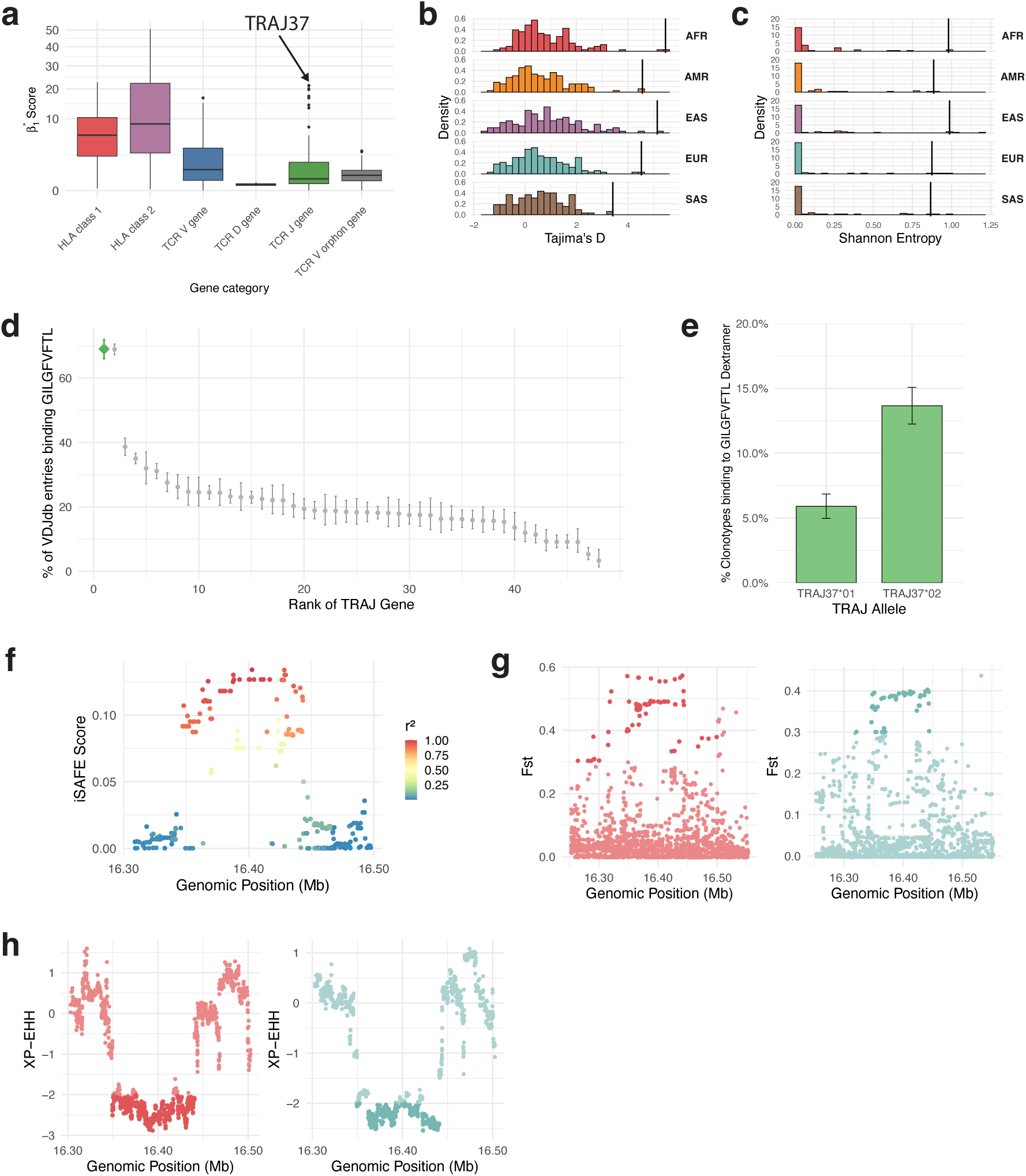
Balancing and positive selection on specific TCR alpha V and J genes. (a) Distribution of BetaScan-computed β_1_* scores of balancing selection across the TCR genes, genes known to be subject to balancing selection (HLA class 1 and 2), and genes known to be under neutral evolution (TCR V orphon genes). (b) Distribution of Tajima’s D across the TRAJ gene array; the vertical line indicates the TRAJ37 gene. (c) Distribution of the Shannon entropy of one-field allele frequencies across TRAJ genes. TRAJ37’s entropy is indicated with vertical line. (d) Percentage of VDJdb entries of each gene binding the GILGFVFTL influenza matrix protein epitope. TRAJ37 is shown in green. (e) Percentage of clonotypes binding to GILGFVFTL dextramer among clones expressing allelic variants of TRAJ37. (f) iSAFE plot to detect variants subject to positive selection in individuals of East Asian ancestry. Color indicates linkage (r^2^) to iSAFE lead variant. (g) Population differentiation (Fst) between East Asian vs African ancestry (left) and East Asian ancestry vs European ancestry (right). Fst values above 0.3 are shaded darker. (h) XP-EHH statistic comparing the East Asian ancestry’s haplotype structure to that of African ancestry (left) and to that of European ancestry (right). XP-EHH values less than – 2 are shaded darker, indicating East Asian donors exhibit longer haplotypes in this region.

Unlike the TCRβ chain’s CDR3, which is primarily shaped by junctional insertions, the TCRα chain’s CDR3 is more strongly determined by its J gene^60^. Thus, we wondered if TRAJ37 was subject to balancing selection because it contributed to the recognition of an immunodominant antigen. To investigate this, we analyzed antigen specificity data from VDJdb^61^. TRAJ37 was strongly enriched in TCRs binding the key GILGFVFTL epitope of the influenza matrix protein (Fisher’s exact p-value < 10^-16^), with nearly 70% of TRAJ37^+^ TCRs in VDJdb binding this peptide (**Fig. 4d**). Prior studies have found TRAJ37 contributes to a dominant TCR clonotype that recognizes this epitope on a common HLA allele (A*02:01), and structural analyses show the CDR3α makes critical contacts with the peptide^62,63^. If balancing selection acted on TRAJ37 to enable recognition of diverse influenza-derived epitopes as the pathogen evolved, alleles of TRAJ37 may be differentially capable of recognizing those epitopes. Indeed, by analyzing a single-cell dataset^64^ of CD8^+^ T cells stained with a influenza GILGFVFTL–A0201 pMHC dextramer (**Methods**), we found TRAJ37*02^+^ cells were significantly more likely to recognize this pMHC than TRAJ37*01^+^ cells (p = 4.3 × 10^-7^; 13.1% vs 6.1%; **Fig. 4e**). These two alleles differ by a nonsynonymous change in the CDR3α region. The retention of multiple TRAJ alleles encoding distinct antigen-contacting residues may have conferred a selective advantage in the face of recurrent exposure to a rapidly-evolving virus.

Additionally, we detected balancing selection in TCR V genes, with the strongest signal in TRAV14/DV4 (**Supplementary Fig. 6b**). Despite these notable examples, variants within TCR loci generally did not show evidence of balancing selection, especially compared to signals observed in the classical HLA genes (**Fig. 4a**). This, in part, may be due to the differences in the genetic architecture of these loci; on a given haplotype there are only three HLA class I genes that present peptides to CD8^+^ T cells, while there are dozens of TCR V and J genes. This may buffer the pressure for balancing selection to act broadly on all TCR genes.

Finally, we asked if positive selection may have acted on specific variants in TCR genes. To investigate this, we used integrated Selection of Allele Favored by Evolution (iSAFE)^65^ analysis to identify variants with evidence of positive selection. The strongest signal emerged from the TRAV gene array in donors of East Asian ancestry (**Fig. 4f**). This region exhibited elevated population differentiation, with Fst values exceeding 0.4, and the lead variant identified by iSAFE had a higher allele frequency in East Asian donors (0.67) relative to African donors (0.10) and European donors (0.19), consistent with selection on standing variation, often termed a ‘soft sweep’ (**Fig. 4g**; **Supplementary Fig. 6c**). To further investigate this signal, we applied haplotype-based statistical methods and site frequency spectrum-based methods. Integrated haplotype score (iHS)^66^ analysis supported a signature of directional selection, with SNPs in this region exhibiting extreme |iHS| > 2, and cross-population extended haplotype homozygosity tests (XP-EHH)^67^ comparing the East Asian haplotype structure in this region to that of African and European ancestries corroborated the population-specific nature of this selective event (Fig 4h; **Supplementary Fig. 6d**). Moreover, Fu and Li’s D* and F* statistics^68^ were markedly negative in this region (< −2), reflecting a skew of the site frequency spectrum consistent with directional selection (**Supplementary Fig. 6e-f**).

Within the haplotype tagged by the iSAFE signal (r² > 0.85), we found two candidate causal variants in coding or regulatory regions of TCR genes: a splice acceptor variant in TRAV26-2 which renders the gene nonfunctional and a framework 3 region nonsynonymous variant in TRAV30, located three amino acids away from the CDR2 loop. Both of these variants are estimated to have arisen prior to out of Africa migration, over 9,200 generations ago.^69^ Due to the tight linkage between these candidates, unambiguously identifying the causal driver variant is a challenge, as it is for many other loci under positive selection. However, because mutations in framework regions have been shown to alter T cell specificity^52,53^, we hypothesize that the TRAV30 framework variant enhanced germline-encoded recognition of a specific pathogen circulating in ancestral East Asian populations, leading to its positive selection.

### Influence of TCR allelic polymorphism on single-cell phenotypes

Having established that polymorphism in TCR genes has been shaped by selective forces, we wondered how this class of genetic variation may influence core functional properties of T cells. Although the TCR is central to T cell identity, the impact of inherited TCR variation on human T cell biology remains largely unexplored^18^. To investigate this, we leveraged the near-complete map we developed of common human TCR alleles to genotype TCR chains in published single-cell datasets and tested how allelic polymorphism influences (1) cellular fate commitment, (2) clonal expansion in disease contexts, and (3) cell surface TCR abundance (**Fig. 5a**). Unlike most single cell association frameworks, our covariate of interest is germline encoded, so identified signals likely represent causal effects mediated by the TCR genotype.

**Figure 5:**
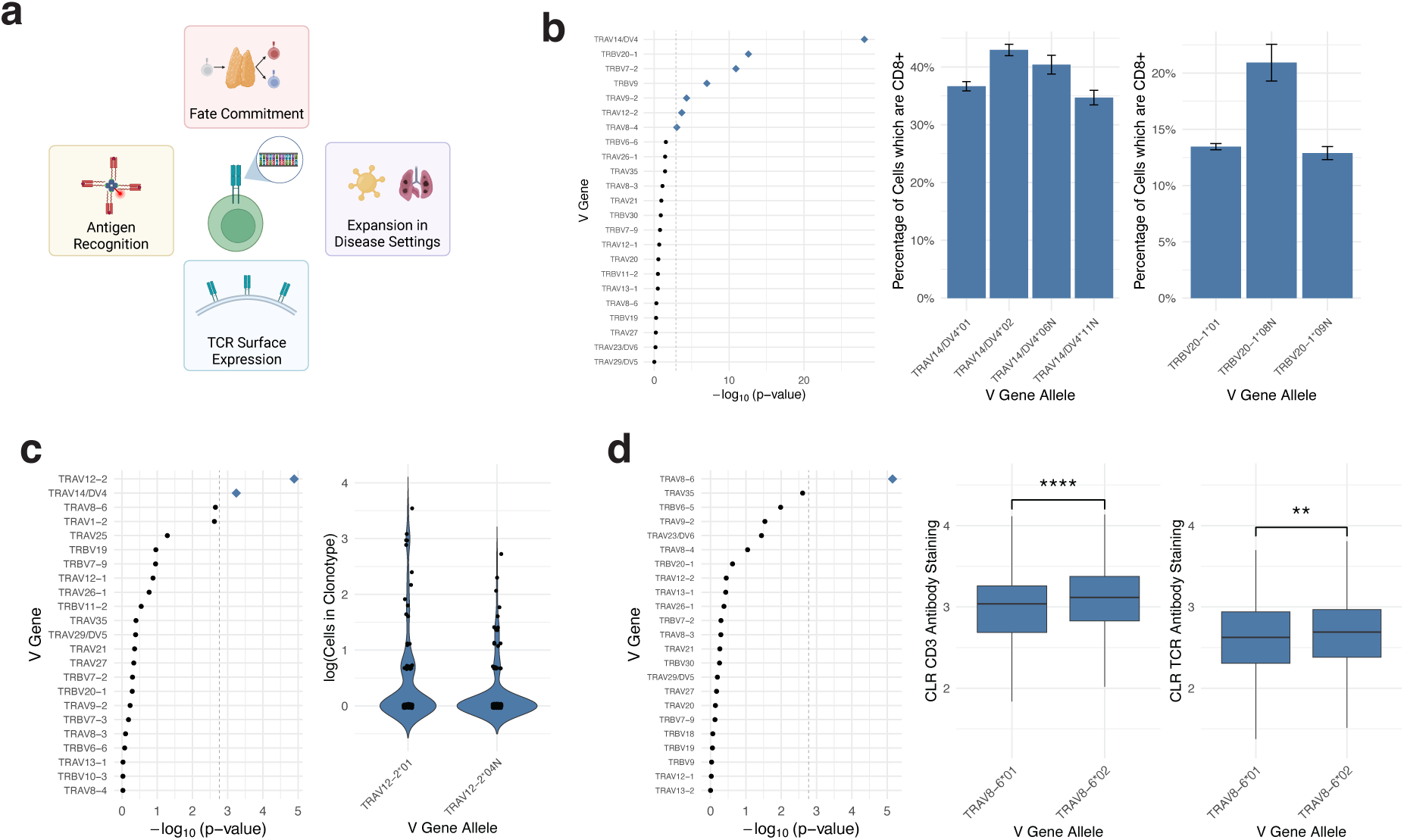
Influence of TCR allelic variants on T cell functional phenotypes. (a) Schematic of the single-cell analytical framework. A population-scale map of TCR alleles was used to genotype individual T cells from single-cell TCR-sequencing datasets to discover how allelic polymorphisms influence cellular phenotypes. (b) Influence of V gene alleles on CD8⁺ fate commitment. Left: Nominal p-value from a likelihood ratio test comparing a null mixed-effects model to a full model including an allele term (i.e., omnibus test of allelic effects). The vertical dotted line indicates the Bonferroni-corrected significance threshold; blue diamonds denote genes with p-values passing this threshold. Right: Proportion of cells adopting a CD8⁺ fate stratified by allele. (c) Effect of V gene allelic variants on clonal expansion of CD8⁺ memory T cells in donors with COVID-19. Left: Nominal p-values from an omnibus mixed-effects linear regression model. Right: Expansion of CD8⁺ memory T cells stratified by allele. (d) Impact of V gene alleles on TCR surface abundance. Left: Nominal p-values from an omnibus mixed-effects linear regression model. Right: Centered log ratio normalized staining of CD3 antibody and pan–TCRαβ antibody of T cells carrying different alleles of TRAV8-6.

Recent studies, including our own, have shown the TCR sequence influences thymic lineage commitment and continues to guide fate decisions in the periphery^70–72^. During thymic selection, for example, the specific V genes a T cell expresses biases its likelihood of CD8 lineage commitment^70,71^. Thus, we hypothesized that germline variation within individual V genes may modulate fate determination and employed an omnibus logistic regression strategy to a combined cohort of 522 donors to test this hypothesis (**Methods**). After controlling for potential donor- and age-mediated effects and randomly sampling one cell per clonotype, we found seven V genes exhibited significant allele-dependent effects on CD8^+^ fate acquisition (**Fig. 5b**). We observed the strongest signal for TRAV14/DV4^+^ cells (omnibus p = 8.5 × 10^-29^); 36.7% of cells expressing TRAV14/DV4*01 were CD8^+^, while 42.9% of cells expressing TRAV14/DV4*02 were CD8^+^. These two alleles differ by nonsynonymous changes in CDR1 and CDR2, regions known to extensively contact pMHC ligands during thymic selection^47^.

Extending this analysis to peripheral fates, we found TCR germline variation also influenced the likelihood of memory formation. Among TRAV9-2^+^ cells, 46.1% of TRAV9-2*02^+^ cells exhibited a memory phenotype while 48.5% of TRAV9-2*11N^+^ cells did (omnibus p = 1.4 × 10^-4^; **Supplementary Fig. 7a**). Collectively, these findings suggest common genetic variation in TCR genes influences both thymic and peripheral T cell fate acquisition.

We next asked whether TCR allelic variation alters T cell responses during disease, first examining SARS-CoV-2 infection. Since clonal expansion of CD8^+^ memory T cells is a hallmark of antiviral responses, we tested whether cells carrying a particular V gene are differentially expanded based on the allele they carry in 88 SARS-COV-2 positive donors^73^. Greater expansion of clones carrying a particular allele may suggest superior viral antigen recognition. We observed the strongest effect in TRAV12-2^+^ clonotypes (omnibus p = 1.29 × 10^-5^); TRAV12-2*01^+^ clonotypes were on average 1.95 times more expanded than TRAV12-2*04N^+^ clonotypes (**Fig. 5c**). Prior structural studies show the TRAV12-encoded CDR1α makes extensive contacts with the immunodominant spike protein derived S269-277 peptide presented on HLA-A*02:01, contributing 44.4% of the α chain’s buried surface area—more than any other region, including CDR3^74^. TRAV12-2*04N harbors a nonsynonymous mutation at IMGT position 29 within the center of the CDR1α; this mutation may reduce affinity to this immunodominant peptide.

Next, to test whether TCR germline variation influences behavior within the tumor microenvironment, we applied our modeling strategy to a dataset of 12 non-small cell lung cancer tumors^75^. TRAJ24^+^ T cells exhibited significant allele-dependent differences in clonal expansion (omnibus p = 9.7 × 10^-5^); TRAJ24*03^+^ clonotypes were on average 1.85 times more expanded than TRAJ24*02^+^ clonotypes (**Supplementary Fig. 7b**).

Beyond influencing antigen specificity, TCR sequence variation may also modulate receptor surface expression, a key determinant of signaling strength. Artificially-introduced framework mutations can alter surface expression^54^, so we asked whether common inherted variation in V genes has a similar effect. To test this, we applied our mixed-effects modeling approach to TCR+CITE-seq data from 122 donors^76^, using normalized anti-CD3 antibody staining a proxy for surface TCR abundance. We focused on CD4^+^ naïve cells, the most abundant subset. We observed the strongest effect in TRAV8-6^+^ cells (omnibus p = 7.05 × 10^-6^). TRAV8-6*02^+^ CD4^+^ naïve cells exhibit significantly increased surface staining of both the CD3 and pan-TCRαβ antibodies in the discovery cohort, an effect that was replicated in an independent cohort profiled with a distinct anti-CD3 antibody (**Fig. 5d**; **Supplementary Fig. 7c**). The two common TRAV8-6 alleles, TRAV8-6*01 and TRAV8-6*02, differ by two residues at IMGT positions 66 and 67 in the framework 3 region. Increased TCR surface expression may enhance sensitivity to cognate pMHC ligands; further experimental work is needed to determine how germline variation influences this sensitivity^54^.

## Discussion

Our study highlights a previously under-characterized class of human genetic diversity within loci central to adaptive immunity and suggests that it has functional implications. Evolutionary analyses reveal that TCR genes bear signatures of three distinct evolutionary forces — diversifying, balancing, and positive selection. Selection only occurs when genetic variants confer interindividual fitness advantages. The fact that natural selection is observed in genes encoding the TCR — a component of the adaptive immune system traditionally thought to function through somatically-generated stochastic diversity — suggests that inherited TCR variation critically shapes immune responses.

Though our approach was equally powered to detect signals in both the TRA and TRB loci, the strongest evolutionary and functional associations consistently localized to TRA. Surprisingly, individual TRAJ genes exhibited greater allelic diversity than TRBJ genes and harbored fewer LoF mutations, even though there are nearly four times as many TRAJ genes as TRBJ genes. This suggests that the two TCR chains may achieve complementary layers of diversity. While the CDR3β region derives much of its diversity from stochastic junctional processes, the CDR3α region is more strongly shaped by inherited V and J genes. Germline polymorphism in TRA genes provides a substrate for natural selection to act upon, enabling populations to retain an immunogenetic “memory” of prior pathogen exposures. This conceptually parallels a recent study finding the kappa and lambda light chains (the immunoglobin analog of the TRA chain) harbor germline-encoded biases towards recognizing common public epitopes of infectious pathogens^77^.

Furthermore, despite the central role of T cells in human immunity, disease associations to the TCR locus remain sparse, with the most notable example being an association between a variant in the TRA V gene array and narcolepsy^78–80^. This gap may well be driven by technical limitations; short-read sequencing fails to accurately genotype the TCR loci, and SNP arrays have poor coverage in these regions^11,13,18^.

Beyond this study, mounting evidence from structural biology, receptor engineering, and repertoire sequencing indicate germline-encoded motifs play key roles in shaping antigen recognition^5–8,30,52–54^. Our ongoing work uses high-quality haplotypes developed from long reads to impute TCR genetic variants within large, biobank-scale cohorts. For many immune-mediated conditions, currently-assayed variants explain a fraction of disease heritability; TCR polymorphisms may contribute to some of this ‘missing heritability’^81^. In certain contexts, HLA risk alleles may act synergistically with TCR risk alleles to modulate genetic risk. More broadly, the overall number of functional TCR alleles an individual carries may also influence disease risk.

Furthermore, we discovered LoF alleles in key autoimmune-associated TCR genes. Determining whether these LoF alleles are protective through genetic association studies would directly inform therapeutic strategies that ablate autoreactive T cell subsets, such as recent efforts targeting TRBV9^+^ cells in ankylosing spondylitis^82^. Furthermore, the commonality of such LoF alleles suggests that V and J genes are not haploinsufficient, supporting the safety of such interventions^83^.

While we focused on common alleles in this study, we observed hundreds of rare one-field alleles present below 0.01 frequency in all ancestries. Rare V gene alleles, in fact, outnumbered common V gene alleles. Given growing evidence that germline residues can mediate autoantigen recognition, it is plausible that these alleles may contribute to rare syndromes of autoimmunity^29,30,84^. Additionally, our map of TCR germline diversity serves as a valuable reference to the immunogenetics community; studies profiling the TCR repertoire can employ this resource to genotype individual TCR chains at allele, rather than gene, resolution.

Our study faced several limitations. Limited TCR sequencing data are available for γδ cells, preventing detailed single-cell analysis of this population. Additionally, the long-read sequencing cohorts in our study did not have matched TCR repertoire data available, limiting our ability to link regulatory variants to repertoire characteristics. Future studies integrating these modalities will be essential to characterize regulatory control of the TCR and quantify the heritability of TCR repertoires.

Ultimately, the TCR repertoire possesses a remarkable ability to recognize a vast universe of antigens. While the somatic variation produced by V(D)J recombination is a powerful mechanism for generating diversity in the repertoire, it does not act alone; it is complemented by extensive germline polymorphism within the TCR loci. Our findings suggest this germline variation shapes interindividual differences in T cell responses, motivating further investigation.

## Methods

### TCR Allele Discovery

Sequencing data were acquired from the Human Pangenome Reference Consortium’s (HPRC) second release and the NIH All of Us CDRv8 release. All samples in the CDRv8 release with PacBio HiFi sequencing data were included, and samples flagged for QC issues or relatedness were removed from consideration.

A custom pipeline was developed to identify TCR alleles from genome assemblies and quantify read support. First, TCR V, D, and J alleles on IMGT-GENEDB (downloaded on December 12, 2024) were aligned to the T2T-CHM13v2.0 reference using Minimap2 (v2.28) to create a custom annotation of the TRA, TRB, and TRG loci on chromosomes 14 and 7. The T2T-CHM13v2.0 reference was chosen over the GRCh38 reference, as the latter reference lacks sequence content for several TRBV genes. Hifiasm was previously applied to build haplotype-resolved de novo assemblies for the HPRC and CDRv8 samples. Contigs from these assemblies were aligned to T2T-CHM13v2.0, and continuous contig segments uniquely mapping to the coordinates of each gene unit (L-V-GENE-UNIT for V genes, D-GENE-UNIT for D genes, and J-GENE-UNIT for J genes) were extracted as candidate allele sequences. HiFi CCS reads of each sample were then mapped back to these candidate allele sequences. An allele was called if at least 8 reads exactly supported the sequence present in the assembly in one donor, or if at least 4 reads exactly supported the assembly sequence across at least 3 donors. The latter condition improved the sensitivity of allelic discovery in the CDRv8 cohort, as many of these samples were sequenced at low depths. This strategy enabled the robust discovery of TCR alleles.

Alleles were discovered at both one-field and two-field resolution. Following IMGT convention, one-field alleles were defined by nucleotide changes in their coding (V-REGION/D-REGION/J-REGION) sequences. For V and D genes, two-field alleles were defined by nucleotide changes in the IMGT-defined GENE-UNIT. For J genes, the gene unit was expanded to include the donor-splice site, which lies directly 3’ of the J-GENE-UNIT. Any allele sequence absent from IMGT-GENEDB was designated as novel and annotated with an ‘N’ suffix. Digger^85^ v0.07.0 was used to annotate the functionality of TCR alleles; it assesses the validity of recombination signal sequences, leader sequences, canonical motifs, and sequence translation. Additionally, genes with non-‘GT’ splice donor sites and non-‘AG’ splice acceptor sites were determined to be nonfunctional.

To orthogonally validate alleles using data from expressed TCR repertoires, FASTQ files from several large single-cell studies that profiled full-length TCR transcripts with the 10x Genomics 5′ V(D)J kit were reprocessed (**Supplementary Table 1**). In additional, all sequencing runs matching the query “(((5’) AND TCR) AND VDJ) AND “Homo sapiens”[orgn: txid9606] NOT (3’) NOT (3 prime)” were downloaded from the NCBI Sequence Read Archive on December 22, 2024. Those found to contain valid TCR sequence content were processed with CellRanger V(D)J 8.0. A BLAST database containing the coding regions of germline TCR alleles observed in our cohort was constructed, and Igblast^86^ v1.22.0 was used to genotype all TCR chains present in the cellranger ‘filtered contigs’ outputs across the datasets. A novel allele was deemed validated in expression data if it was found in at least five cells with a 100% sequence identity match against the expressed chain (as determined by Igblast) and no ties.

To estimate the theoretical number of common circulating alleles, a Michaelis-Menten curve, which has been extensively used to estimate species richness^87^, was fit to the rarefaction curve of each ancestry; the estimated asymptote was taken to be the theoretical number of common alleles circulating in the population.

For the analysis of per-donor functional allele counts, we included only HPRC samples whose genome assemblies fully spanned both the TRA and TRB loci. The number of distinct functional one-field alleles was computed for every donor; each functional allele was counted once, regardless of zygosity. To partition the variance in interindividual functional allele counts, we fit an ordinary least squares linear regression model and performed a Shapley decomposition of model R^2^. In addition, graphs showing the frequency of specific nonfunctional alleles represent the frequency of these alleles computed within the HPRC cohort.

To compute the Shannon entropy of each TCR gene, we employed the following formula: − ∑*_x_*_∈x_ *p*(*x*) *log*_2_ *p*(*x*), where *X* represents the set of alleles for that gene, *x* represents a specific allele of that gene, and *p*(*x*) represents the frequency of that allele.

In all boxplots throughout the manuscript, the bounds of the box represent the first and third quartiles (Q1 and Q3), the center represents the median and the whiskers extend from Q1 − 1.5 × interquartile range to Q3 + 1.5 × interquartile range.

### Genetics Analyses

For evolutionary genetics analyses, we focused on generating and analyzing a joint variant callset from the All of Us CDRv8 release, as sample preparation, sequencing, and bioinformatic processing were done in a uniform manner across this cohort. Per-sample variant calling was previously performed with DeepVariant against the T2T-CHM13v2.0 reference, and samples flagged for QC issues or relatedness were removed from consideration. Per-sample variant calls for chromosomes 7, 9, and 14 were jointly called with GLNexus^88^ v1.4.1 and underwent quality control by filtering out multiallelic sites and variants with minor allele frequency < 0.01, missingness > 0.1, or QUAL < 30. SHAPEIT4^89^ was used to phase the resulting joint callset.

To determine codon-level diversifying selection, Fast Unconstrained Bayesian AppRoximation^45^ and Fixed Effect Likelihood^46^ methods were run as implemented on Hyphy^90^ v2.5. The R seqlogo package v3.2.1 was employed to visualize the amino acid alignments, and the R ape package v5.8.1 was used to plot radial phylogenetic trees.

For analysis of balancing selection, BetaScan^57^ was run with default parameters to compute the β_1_* statistic across all loci considered. To determine which ligands TRAJ37 may bind to, antigen specificity data was downloaded from VDJdb^61^ on April 20^th^, 2025. VDJdb entries from 10x dCODE dextramer experiments were removed. A Fisher’s exact test was used to quantify the enrichment of TRAJ37 in specific antigen binding groups.

To identify candidates of positive selection, iSAFE^65^ v1.0.4 was applied with the following parameters: MaxFreq = 0.8 and MaxRank = 300. Furthermore, selscan^91^ v1.2.0a was employed to calculate and normalize the haplotype-based statistics across chromosome 14. The R popgenome^92^ package v2.7.5 was used to compute Tajima’s D and Fu and Li’s D* and F*. Fst was computed using the Weir and Cockerham method implemented in vcftools v0.1.14.

### Single-Cell Analyses

To test for associations between TCR V and J gene alleles and single-cell phenotypes, we employed mixed-effects regression models implemented in R lme4 package v1.1.23. For all models, only cells with a single TCRα chain and a single TCRβ chain and whose V and J genes could be confidently genotyped (with an IgBlast sequence identity of 100% and no ties) were considered. Additionally, for all models, a single cell was selected at random to represent each expanded clone, as cells from the same clone share identical TCR sequence features and similar transcriptional fates, so do not represent independent observations.

An omnibus regression framework was employed to evaluate the influence of TCR allelic polymorphism on cellular phenotypes. The models were formulated as follows:

Null Reduced Model:

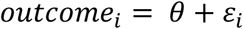

Full Model:

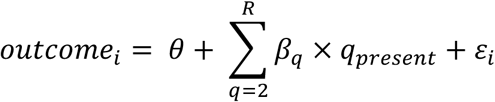

Each observation *i* represents a T cell from a donor *d*. The models were run once per TCR gene, where *θ* is the intercept. In the full model, *β_q_* is a fixed effect representing the effect of allele *q* on the outcome, *q_present_*, is an indicator variable with a value of 1 if that allele was observed in the cell, and *R* is the number of alleles observed for that gene. A likelihood ratio test was used to compare the fit of the null reduced model to that of the full model. For each cellular phenotype, additional covariates were included in both the reduced and full models, as described below:

To test for V gene allele influences on fate commitment, an omnibus mixed effects logistic regression approach of the above form was run on a combined cohort of 522 donors from three recently-published 5’ single-cell datasets with TCR-sequencing information^73,76,93^. An additional fixed-effect for donor age, a fixed effect for donor sex, a fixed effect with three levels representing the dataset the cell was observed in, a fixed effect for donor COVID-19 status, and a random effect intercept for each donor was included. For the model of CD8^+^ fate commitment, outcome variable was set to 1 if the cell was in a CD8^+^ transcriptional cluster, and 0 if the cell was in CD4^+^ transcriptional cluster based on the cell annotations provided by the authors. For the model of memory commitment, the outcome variable was set to 1 if the cell was in a memory cluster, and 0 if the cell was in a naïve cluster. For both outcomes, all non-CD4^+^/CD8^+^ T cells were excluded. This model was run once per TRAV and TRBV gene; V genes with at least two alleles expressed in at least 5000 clonotypes were considered.

To test for V gene influences on antigen binding, an omnibus fixed effects logistic regression approach of the above form to was applied to dataset of CD8^+^ T cells from four donors profiled using 10x genomics dextramer staining technology^64^. Dextramer binding calls were computed using our previously-described approach, wherein a negative binomial regression model was employed to remove background staining and strict thresholds were set to distinguish binders from non-binders; this strategy regresses out TCR expression and surface staining to isolate effects due to variation TCR sequence content rather than differences in TCR expression^70^. The outcome variable was a binary indicator representing whether or not the cell bound to the influenza GILGFVFTL dextramer. An additional fixed effect was included for each donor.

To test for V gene allele influences on in vivo clonal expansion during infection, an omnibus mixed effects linear regression approach of the above form was fit to a cohort of 88 donors deeply profiled during COVID-19 infection^73^. The outcome variable was the log count of cells observed in the clone. A fixed effect for donor age, a fixed effect for donor sex, and a random effect intercept for each donor was included. All cells in the CD8^+^ memory compartment were considered, the model was run once per V gene, across all V genes with at least two alleles present in at least 200 clonotypes.

To test for allelic influences on clonal expansion in the tumor microenvironment, an omnibus mixed effects linear regression approach of the above form was fit to a cohort of 12 donors whose non-small cell lung cancer tumors were profiled^75^. The outcome variable was the log count of cells observed in the clone. A random effect intercept for each donor was included. The model was run once per V gene, across all V genes with at least two alleles present in at least 200 clonotypes.

Finally, to test for V gene influences on cell surface TCR abundance, an omnibus mixed-effect linear regression model approach with the above form was fit to a cohort of donors profiled with CITE-seq^76^. A fixed effect for donor age, a fixed effect for donor sex, a fixed effect for donor COVID-19 status, and a random effect intercept for each donor was included in the model. CITE-seq staining counts were centered log-ratio normalized (CLR) across all antibody derived tags for each cell. The outcome variable was the CLR-normalized value of the CD3 antibody. All cells in the CD4^+^ naive compartment were considered, the model was run once per V gene, across all V genes with at least two alleles present in at least 1000 clonotypes. The observed TRAV8-6 signal was replicated in a distinct cohort^93^ profiled with a different anti-CD3 antibody.

For all the above models, Bonferroni correction was applied to correct for multiple testing.

## Acknowledgements

We are grateful to the participants in the All of Us study for their contributions, and we would like to thank the National Institute of Health’s All of Us Research Program for making available the participant data examined in this study. We would like to acknowledge the Human Pangenome Reference Consortium (BioProject ID: PRJNA730823) and its funder, the National Human Genome Research Institute (NHGRI). We thank members of the Raychaudhuri lab for helpful discussions. We thank Ryuya Edahiro and Yukinori Okada for sharing single-cell TCR expression data. This work is supported in part by funding from the NIH, including R01AR063759, P01AI148102, U01HG012009, R56HG013083, and T32GM144273.

## Competing Interests

We declare no conflict of interest for this study. S.R. is a founder for Mestag, Inc, and serves as a scientific advisor for BMS, Jannsen, Merck, Pfizer, Nimbus, and Bonito.

## Resource Availability

All sequencing data analyzed in this study were previously deposited in publicly accessible databases. The genetic data analyzed was provided by the All of Us and Human Pangenome Reference Consortium datasets, while the single-cell TCR repertoire expression data was acquired from the published studies listed in **Supplementary Table 1**. Sequences of common validated TCR alleles are available at https://github.com/SreekarMantena/tcrdiversity.

## Author Contributions

S.M. and S.R. conceptualized the project. S.M. performed analyses with guidance from S.R. and A.A. All authors contributed to the final manuscript.

