## Supplementary Material for "TCR germline diversity reveals evidence of natural selection on variable and joining alpha chain genes"

### Supplementary Figures

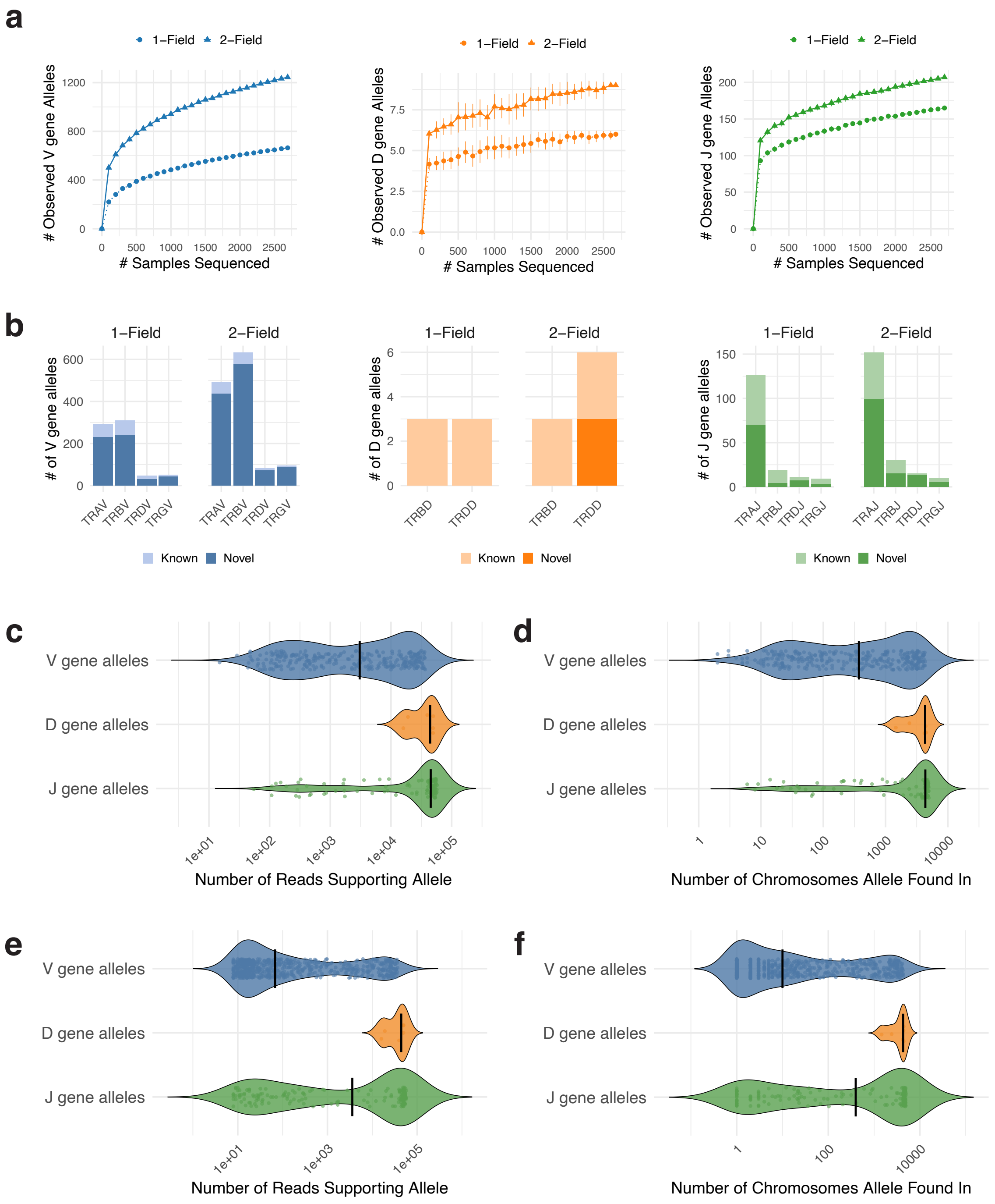


**Supplementary Figure 1 — Allelic discovery of TCR genes. (a)** Rarefaction curves for all V (left), D (center), and J (right) gene alleles, including rare alleles. **(b)** Total number of known and newly discovered alleles for TCR V, D, and J gene groups, including rare alleles. **(c)** Number of reads exactly supporting the sequences of common V, D and J gene alleles. The vertical black line represents the median. **(d)** Number of chromosomes common V, D, and J gene alleles were found in. **(e)** Number of reads exactly supporting the sequences of all V, D and J gene alleles, including rare alleles. **(f)** Number of chromosomes V, D, and J gene alleles were found in, including rare alleles.

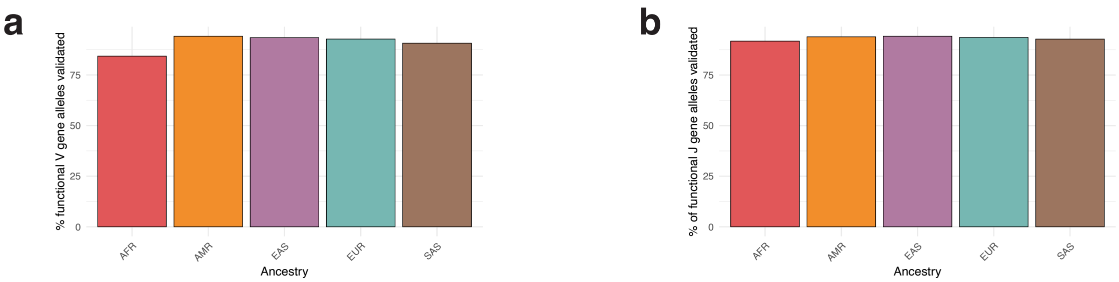


**Supplementary Figure 2 — TCR allele validation in expressed repertoires. (a)** Percentage of functional V genes that are common in each ancestry that were able to be validated in expressed repertoires (see Supplementary Note 1). **(b)** Percentage of functional J genes that are common in each ancestry that were able to be validated in expressed repertoires.

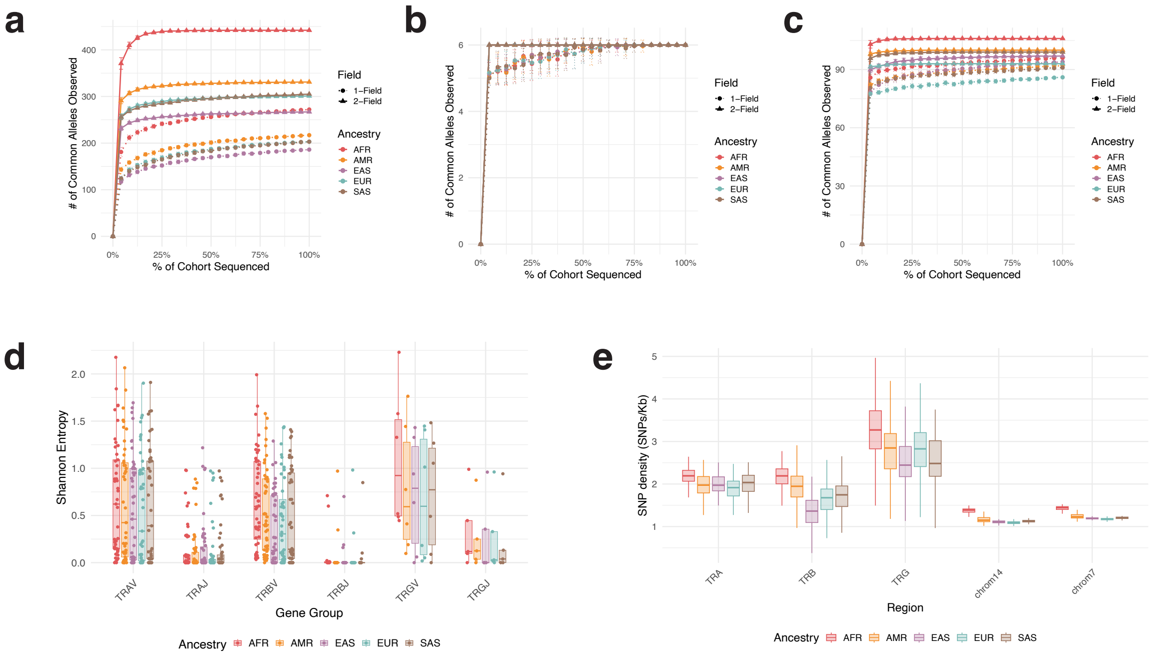


**Supplementary Figure 3 — TCR alleles across ancestries.** **(a)** Rarefaction curves for each of the five ancestries depicting the number of common V gene alleles discovered. **(b)** Analogous to (a), but for common D gene alleles. **(c)** Analogous to (a), but for common J gene alleles. **(d)** Shannon entropy of allele frequencies across TCR gene groups. Each dot represents the Shannon entropy of the one-field allele frequencies of a specific TCR gene in a specific ancestry. **(e)** SNP density by ancestry across the TCR loci and the chromosomes they reside on.

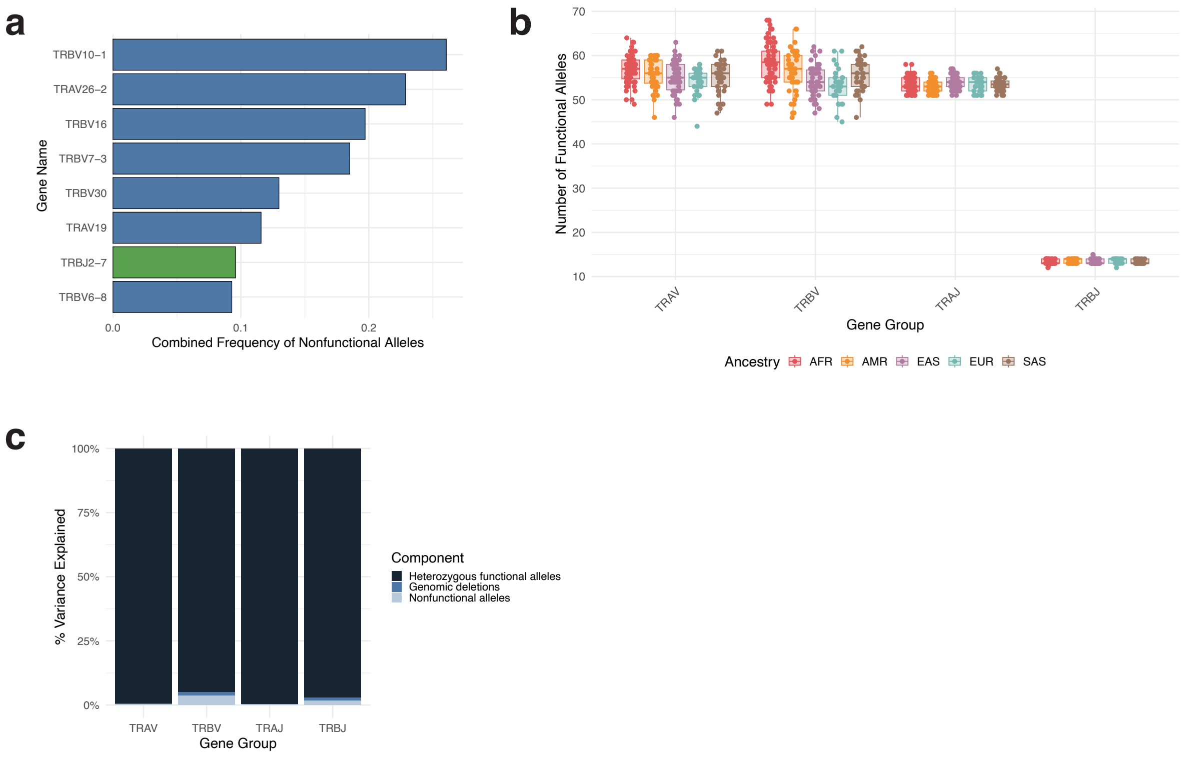


**Supplementary Figure 4 — TCR allele functionality. (a)** TCR V and J genes with a high burden of nonfunctional alleles. **(b)** Distribution of the number of functional TCR V and J genes across donors of different ancestries. **(c)** Variance in the number of distinct functional TCR V and J alleles per individual, partitioned into three components: functional heterozygosity, genomic deletions, and nonfunctional alleles.

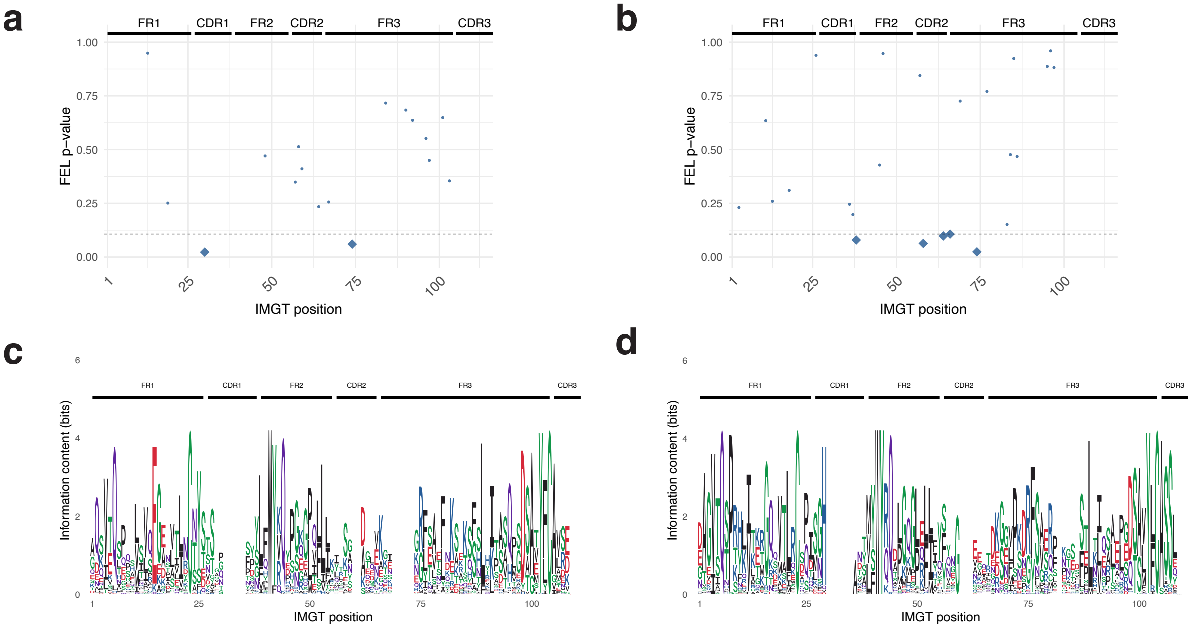


**Supplementary Figure 5 — Codon-level selection on V genes.** **(a)** Results of fixed effect likelihood test (FEL) for diversifying selection on common functional TRAV gene alleles. FEL employs a likelihood ratio test to determine if the nonsynonymous substitution rate is significantly different than the synonymous rate. The p-values from this likelihood ratio test are shown on the y axis; p-values below the dotted line threshold indicate a site is undergoing diversifying selection. Points for positions at which the estimated nonsynonymous substitution rate is greater than the estimated synonymous substitution rate are shown. **(b)** Analogous to **(c)**, but for common functional TRBV gene alleles. **(c)** Amino acid sequence logo for common functional TRAV gene alleles. Sites with no amino acid present represent gapped positions. **(d)** Analogous to (a), but for common functional TRBV gene alleles.

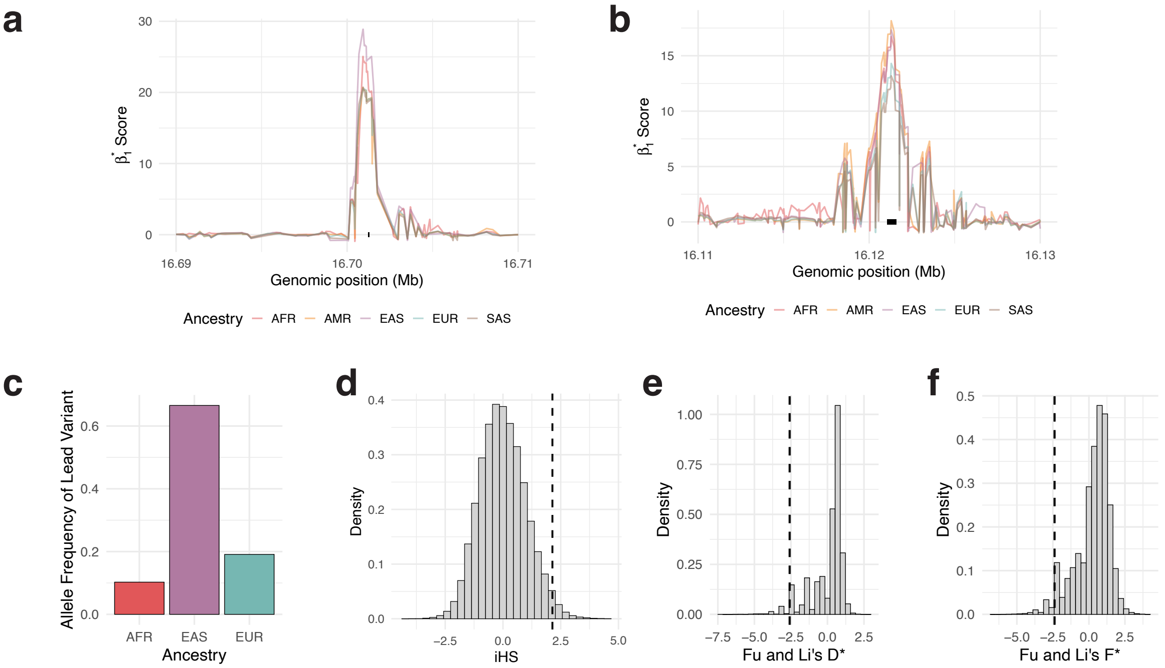


**Supplementary Figure 6 — Balancing and positive selection on TCR genes. (a)** $\beta_1^*$ statistic of region surrounding TRAJ37 gene across ancestries. Gene body of TRAJ37 is indicated with horizontal black bar. **(b)** Analogous to (a), but for TRAV14/DV4. **(c)** Allele frequency of iSAFE lead variant across ancestries **(d-f)** Distribution of iHS (d), Fu and Li's D* (e), and Fu and Li's F* (f) across chromosome 14 in donors of East Asian ancestry, with TRA region under directional selection marked with vertical black line.

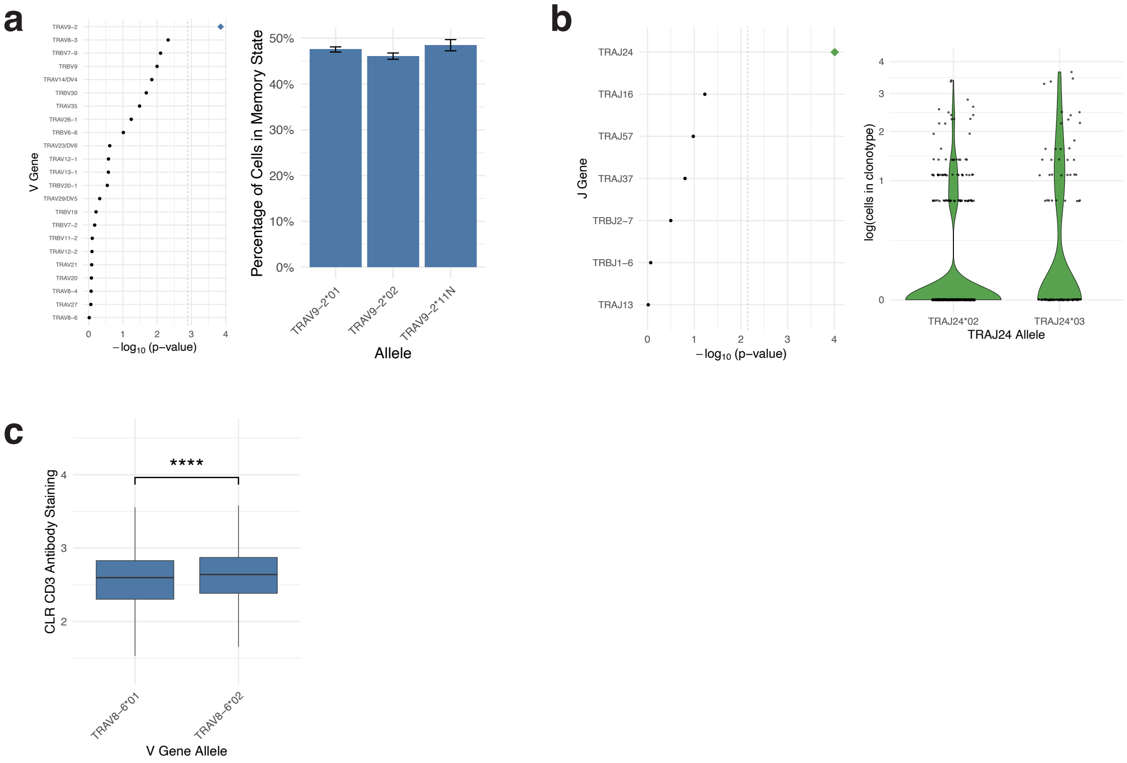


**Supplementary Figure 7 — Allelic influences on cellular phenotypes.** **(a)** Effect of V gene allelic variants on the acquisition of memory fate. Left: Nominal p-values from an omnibus mixed-effects linear regression model. Right: Proportion of TRAV9-2$^+$ cells observed in a memory fate stratified by allele. **(b)** Influence of J gene allelic variants on T cell clonal expansion in non-small cell lung cancer tumors. Left: Nominal p-values from an omnibus mixed-effects linear regression model. Right: Expansion of tumor-infiltrating TRAJ24$^+$ T cells stratified by allele. **(c)** Centered log ratio normalized CD3 antibody staining of TRAV8-6$^+$ cells in validation cohort, stratified by allele.

### Supplementary Note 1

As described in the Methods, to orthogonally validate TCR alleles, we collected several large, publicly available human cohorts profiled with 10x genomics 5' TCR-sequencing technology. In addition, we collected and processed .fastq files 10x 5' TCR sequencing experiments from the NCBI sequence read archive. The total list of studies and sequence read archive runs used for validation is available in Supplementary Table 1. We focused on validating functional alleles for the TRA and TRB chains, as publicly available sequencing data for $\gamma\delta$ T cells remains quite limited.

93.7% of functional V gene alleles common in donors of European ancestry could be validated in expression data, and 93.6% of functional J gene alleles common in donors of European ancestry could be validated (Supplementary Fig. 2). The lowest validation rates were observed for alleles common in donors of African ancestry, with 84.3% of functional V gene alleles and 91.8% of functional J gene alleles common in this ancestry being validated (Supplementary Fig. 2). We suspect this is due to the limited representation of donors of African ancestry within publicly available single-cell sequencing cohorts; indeed, the largest cohorts in our validation set included predominantly donors of European and East Asian ancestry.
